# How does mixed reality affect quiet stance?

**DOI:** 10.1101/129239

**Authors:** Gaiqing Kong, Kunlin Wei, Konrad P. Kording

## Abstract

Mixed reality (MR) has promise for learning, design, and entertainment, and for use during everyday life. However, when interacting with objects in mixed reality, will moving objects make us fall or perturb our postural stability? To address this question, we recruited participants, instructed them to stand quietly, and measured how much virtual objects presented in mixed reality (Microsoft HoloLens) affected their stance. We analyzed the effects of solid object and text, in both a static and a dynamic setting. Mixed reality events induced some movements, but the effect, while significant, was exceptionally small (< 1mm & < 0.5° perturbations in terms of mean distance and angle rotations). We conclude that induced movement in “real reality” should not be too much of a concern when designing mixed reality applications.

## Introduction

Mixed reality promises to improve our interactions with both real and artificial worlds by allowing the two to mix. For example, the senior author on this paper would love a system that can project the name over the head of every person he meets. While mixed reality has the potential to improve education, entertainment, and communication, it requires users to wear a head mounted display (HMD, e.g. Microsoft Hololens) for displaying virtual objects. For safety reason, it is important to study how this technology affects user’s posture and balance. However, the effects of mixed reality using HMD on postural stability and balance is not well understood.

Virtual reality is known to have an influence on posture. Various studies have looked at the effect of virtual reality (VR) on postural stability [1–9]. Earlier studies found that sensory inputs from visual system conflicted with that from vestibular and somatosensory systems induced by VR caused more postural instability [2, 9]. VR also influences body sway and instability [4], producing increased postural sway that is comparable to those observed with closed eyes [1]. Besides its effect on static stance, VR also affects dynamic postural balance [6]. Previous studies have found that displacement of center of pressure or angular deviations of shoulder caused by VR amount to a few centimeters or degrees [1, 6, 9]. One report even documented that standing participants sometimes had to take an extra step to prevent falling when facing with sudden perturbation in VR [7]. These findings in VR settings thus suggest that it is important to quantify the effect of mixed reality on postural stability.

Posture control is complex as the nervous system needs to deal with the information from the body, e.g. vestibulospinal system, proprioception, visual information, as well as information from the environment, e.g. the stability of support and the field of view [10, 11]. Studies on postural stability measurement include ground reaction force [4–6], head movements [7], and upper body movements [1]. However, all measured variables tend to correlate heavily. For instance, the movements of head and upper body directly influence the ground reaction force, though their relationship is not well documented. Here we use head movement only to examine the postural stability in mixed reality.

There are multiple reasons why we may expect an influence of mixed reality on quiet stance. Mixed reality involves stereoscopic visual stimuli that move in the visual field in predictable or unpredictable way. On the other hand, quiet stance is continuously modulated by visual information [10, 12–16]. Furthermore, a sudden appearance of a visual stimulus may elicit a startle effect [17–19] which could lead to quick postural adjustments, including avoidance behaviors [18]. Lastly, a virtual object may affect the visibility of real objects in the external world which typically serve as important reference for maintaining postural stability; this reduction in visibility might affect quiet stance as a result [20]. As such, we expect mixed reality to affect real world stance though estimation of its effect is still lacking.

Here we investigate how static and dynamic objects and text presented in mixed reality affect head movement using visual stimuli generated with the Microsoft HoloLens. We recruited human participants, instructed them to stand quietly and measured the movements of their head using the built-in motion tracking system of the HoloLens. We find extremely small effects of our visual perturbations.

## Method

### Participants

We recruited a total of 22 participants (8 females and 14 males, age: 30.1 ± 7.5, average ± SD). All participants were right-handed, had normal or corrected-to-normal vision without known history of psychiatric or neurological disorders. All participants gave informed consent according to the guidelines of the Institutional Review Board of Northwestern University Medical School to participate in this study.

### Mixed reality environment

Participants stood in the experimental room and wore a head-mounted display (HMD, Microsoft HoloLens, Fig.1). The HoloLens was equipped with a pair of mixed reality smartglasses. It ran on the Windows Holographic platform under the Windows 10 operating system. Hololens features an energy-efficient depth camera with a 120°x120°angle of view. The ambient lighting condition in the experimental room is not too dark and not too bright. The calibration process was performed every day. During the calibration process, the experimenter was asked to align their finger with a series of six targets per eye. It allows the device to adjust hologram display according to the user’s interpupillary distance. The visualizations were first developed in the unity platform (Unity 5.5.0f3 Personal), then we exported the project from Unity to Visual Studio (Microsoft Visual Studio 2015), then built and deployed to the HoloLens. For all the settings, including developing scene in Unity, compiling to Visual Studio, and deploying to Hololens, please refer https://developer.microsoft.com/en-us/windows/mixed-reality/holograms_100. The ‘Near Clip Plane’ of ‘Main Camera’ in our experiment is 0.3. The experimental data was sent to a local sever, and the data analysis was conducted offline by customized Matlab programs (2012b, Mathworks, Natica, MA).

**Fig. 1.**
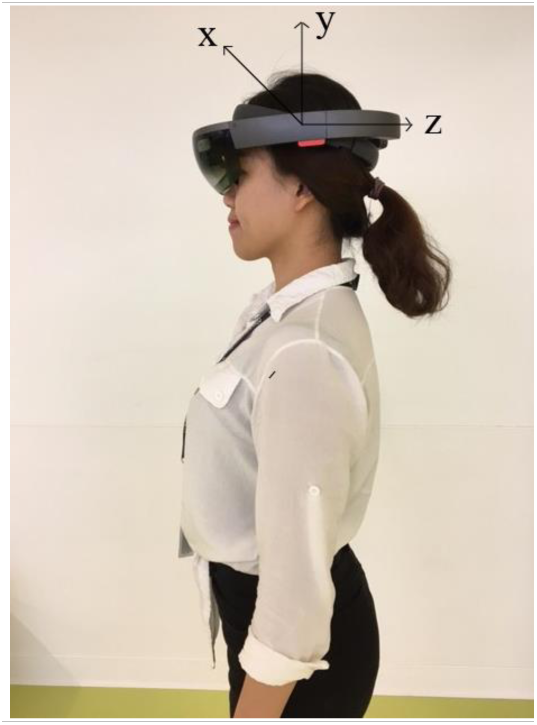
One of the authors testing the study set-up

### Study design

In this experiment, participants were asked to stand comfortably and naturally with the feet parallel and shoulder-width apart and with their arms on their sides. Participants wore the HoloLens and were asked to maintain an upright standing position, remain as stable as possible for the duration of each trial. To obtain identical postural configurations between trials, markings were placed on the ground to guide the placement of the feet of each participant.

During quiet stance, participants viewed four possible visual events, one at a time. These events involved solid object or text, either presented in a static or dynamic setting (Fig. 2). Thus, the experiment had a 2 (object, text) × 2 (static, dynamic) design. The static object was a basketball with a diameter of ∼0.4m appearing 2 meters in front of participants; The dynamic object was the identical basketball moving as 1.8 m/s from 2 meters away toward participants’ face and stopping at 0.2 meters away; The static text was one sentence written as “What does a neuroscientist order at a bar? A spiked drink”, the dynamic text was the same sentence moving as 0.8 m/s from 0.4 meters above to −0.4 meters below and 2 meters in front of participants. For the dynamic settings, the basketball moved in the depth direction towards the participant and the text moved in the vertical direction from high to low. Participants can see all the visual events without moving their heads. In addition, we set a blank screen between every two trials as a baseline condition. The blank screen and the background of the other four visual events in the Hololens are transparent, so what we presented can be mixed with reality. The experiment included 120 trials with each visual event being repeated 30 times. The trials were presented in random order. Before these formal trials, participants practiced for 4 trials where each visual event was randomly present once. Each trial lasted one second, the inter-trial interval was also one second, and the experiment lasted approximately somewhat longer than four minutes. The sampling rate was 40 Hz.

**Fig. 2.**
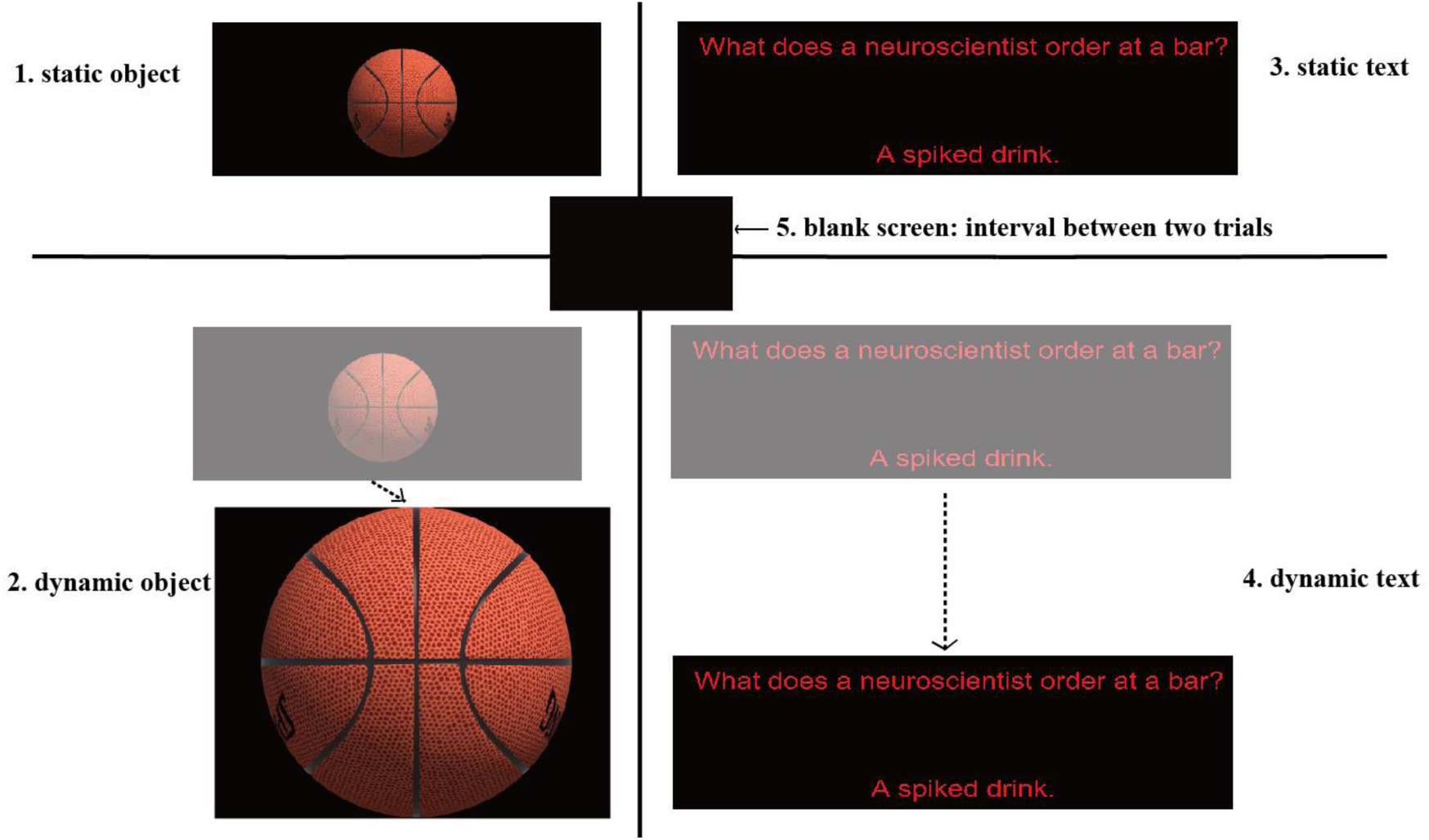
Visual events presented in the Hololens. During quiet stance, participants saw one of four visual objects: solid object (basketball) and text, either present in a static or a dynamic setting. For the dynamic setting, a basketball moved in the depth direction towards the participant and the text moved in the vertical direction from high to low.

### Data analysis

Head sway was measured using the HoloLens system which continuously recorded head position in *x*, *y*, and *z* direction and the corresponding rotation angles (see Fig. 1). We separately analyzed the effects of solid object and text for the static and the dynamic settings. Each trial was normalized by subtracting the beginning state from position and angle variables. We calculated the mean head movements, standard deviation within trials and Root mean square (RMS) values across trials. One-way ANOVAs were performed for each variable. When appropriate, post-hoc comparisons were made between conditions with Bonferroni correction. All statistical analyses were executed using SPSS statistical package (IBM, SPSS 20.0). The significance level was set at α = 0.05.

### Code Availability

We used a set of scripts to control actions of visual events, collect data and send raw data from client to server. All the scripts are made available online at https://github.com/KordingLab/mixed-reality/tree/master. We hope that this code-based will support other scientists and simplify their work with HoloLens.

## Results

To ask how mixed reality affects quiet stance, we recruited 22 healthy participants and instructed them to stand quietly while presenting stimuli in mixed reality (Microsoft Hololens). We measured the head movements using the built-in tracking mechanism of the Hololens (see Fig.1). We presented two classes of stimuli, solid objects vs text, in two different scenarios, static vs dynamic. Averaging across trials allowed us to quantify the effect of mixed reality stimulation onto quiet stance.

As we rely on the measurements from HoloLens, we need to calibrate the HoloLens to ensure that it’s measurements are meaningful and of high quality. We tested 4 positions every 30 centimeters in z direction which we measured using a good old-fashioned ruler. We found a linear relationship between the display measured by Hololens and the actual distance (r = 1.0, *p* < 0.00001), the regression slope is 1 (Fig. 3). We conclude that the measurements by the Hololens camera are reasonably precise.

**Fig. 3.**
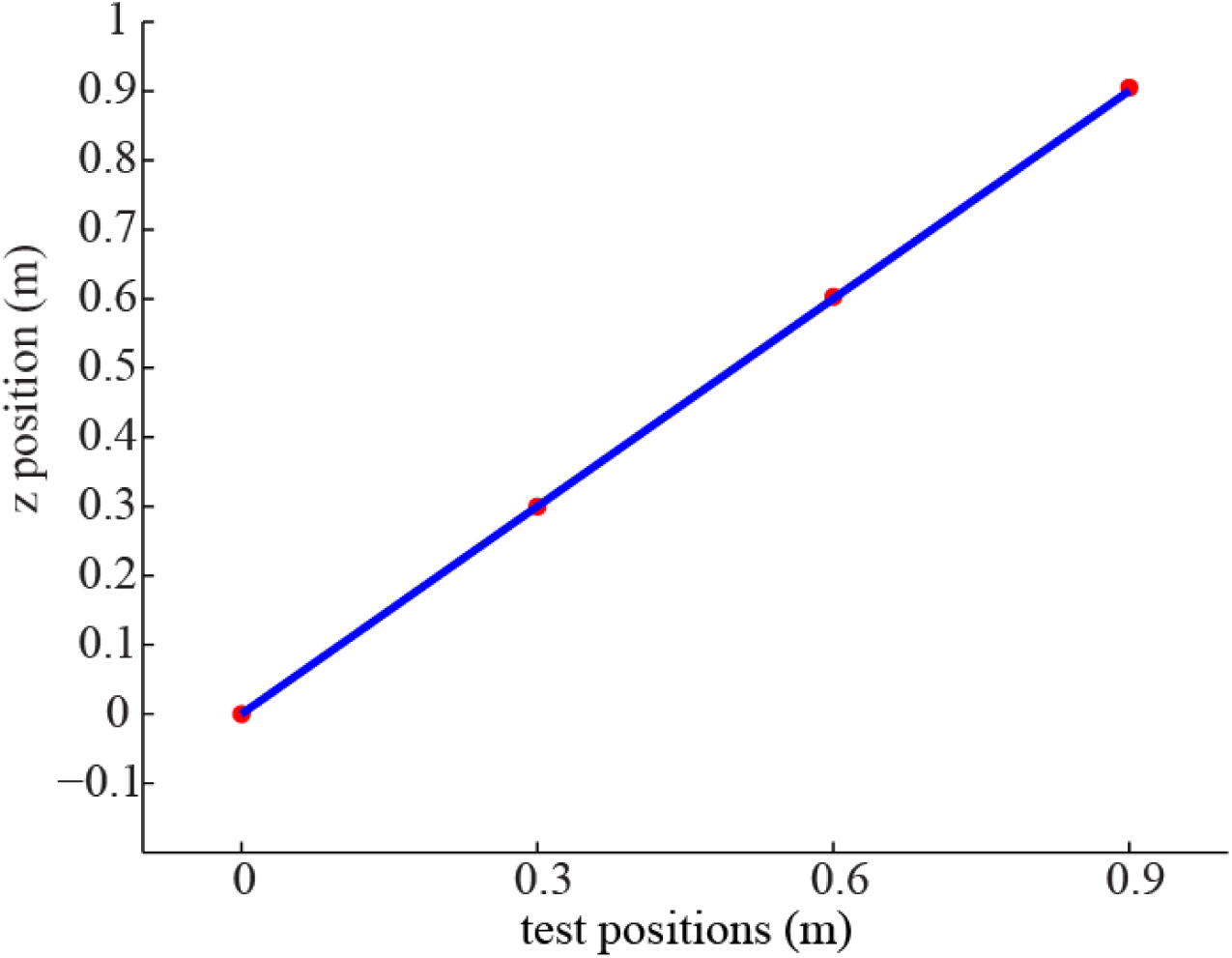
Comparing the actual distance and the display of participant while using Hololens. Four positions every 30 centimeters were tested in z direction. The changes measured by Hololens were very close to the actual distance (r = 1.0), the regression slope is 1. Deviations are probably mostly, due to our inability to precisely position the Hololens by hand for this calibration.

We first want to know if the perturbation induces a meaningful change in the raw position data. To do so we looked at individual trials and looked for movement following the stimulation (Fig. 4). We chose one of the participants and plotted the 10^th^ trial of each kind of visual events. In the single trials that we looked at by eye, there is some ongoing shifts of the body related to the stimulation. An effect may exist, but anecdotally it seems to be very small.

**Fig. 4.**
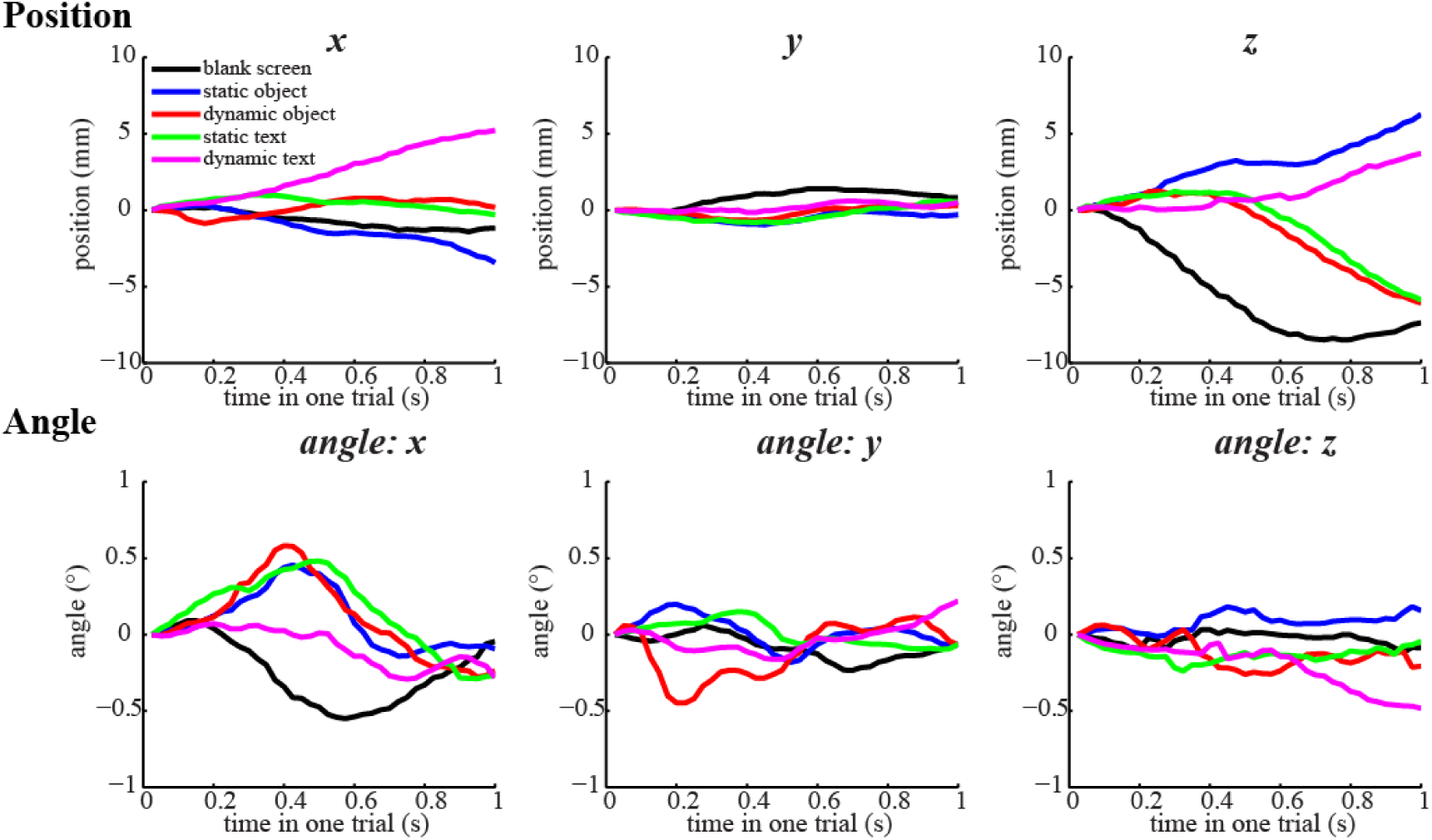
Head movements of single trials during five different visual events – individual participant. Five different visual events were presented to each participant. Position (first row) and angle rotations (second row) were measured and shown one participant. Notice that if angle > 0 participant rotates his/her head counterclockwise around axis (from origin to the axis’ positive direction).

To be able to quantify these small effects we can average across all trials of the same kind. Indeed, when looking at a single participant there seems to be a slight, average influence of the stimulation onto the head movement (Fig. 5). However, the effects are so small or inconsistent that even compiling all data from a single participant we are poorly powered to see the effect.

**Fig. 5.**
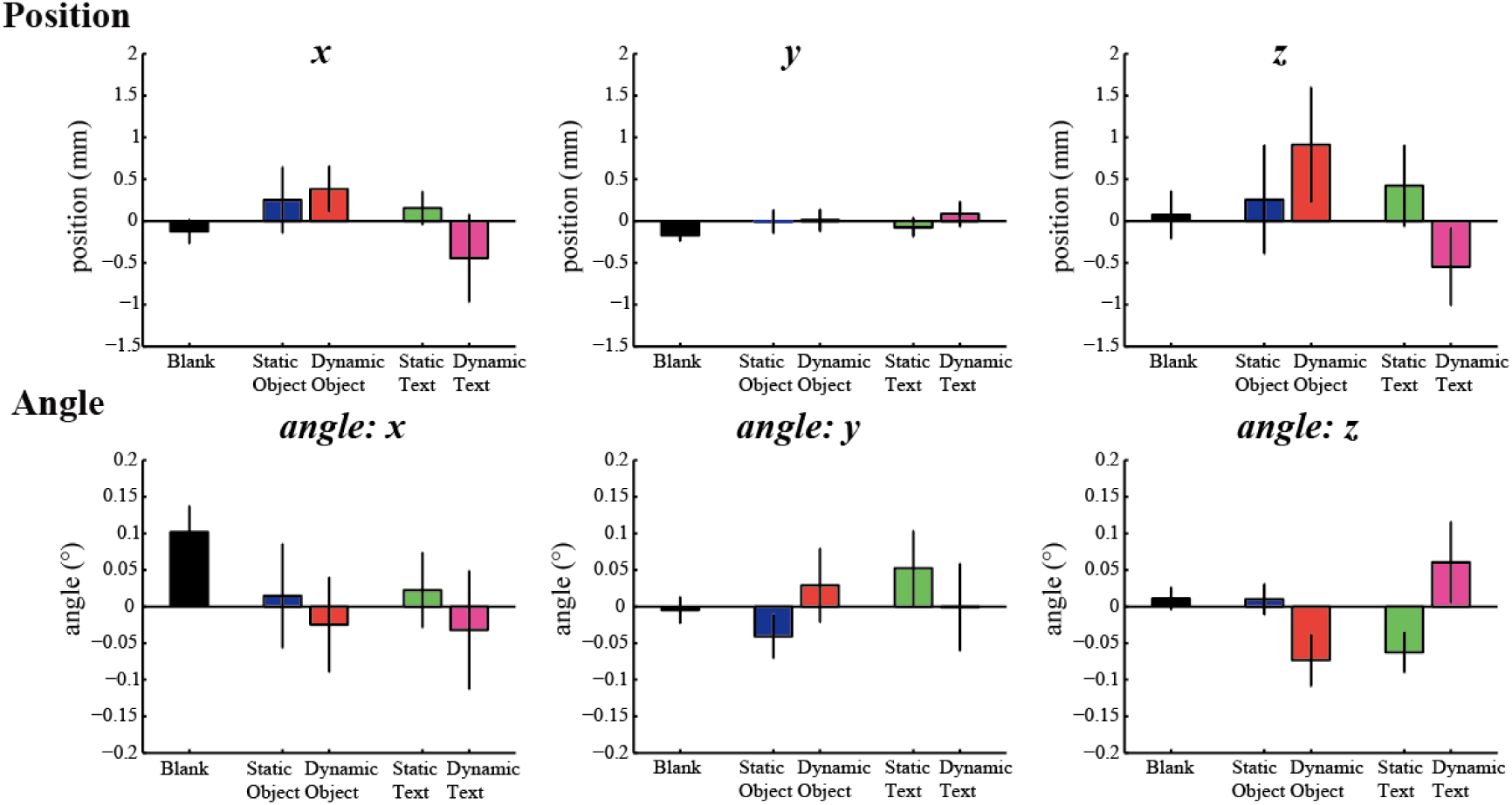
The average distance and angle rotation of head movements across trials from a typical participant. The mean distance (first row) and angle rotations (second row) are shown for five different visual events. Each kind of visual event was randomly presented 30 times. Vertical bars on the columns depict standard error of the mean across trials. Notice that if angle > 0 participant rotates his/her head counterclockwise around the axis (from origin to the axis’ positive direction).

To fully use our statistical power, we can average the data across all participants. Clearly there is an effect in most conditions but it also clearly is extremely small (Fig. 6). The difference between static and dynamic visual events is inconspicuous. It seems that there is an adaptation phase during the first few trials for each kind of visual events but that behavior stabilizes rapidly. All mean perturbations of stance are less than 1cm and 0.5°, even the first ones. But most importantly, anything we see is so small that it is practically irrelevant.

**Fig. 6.**
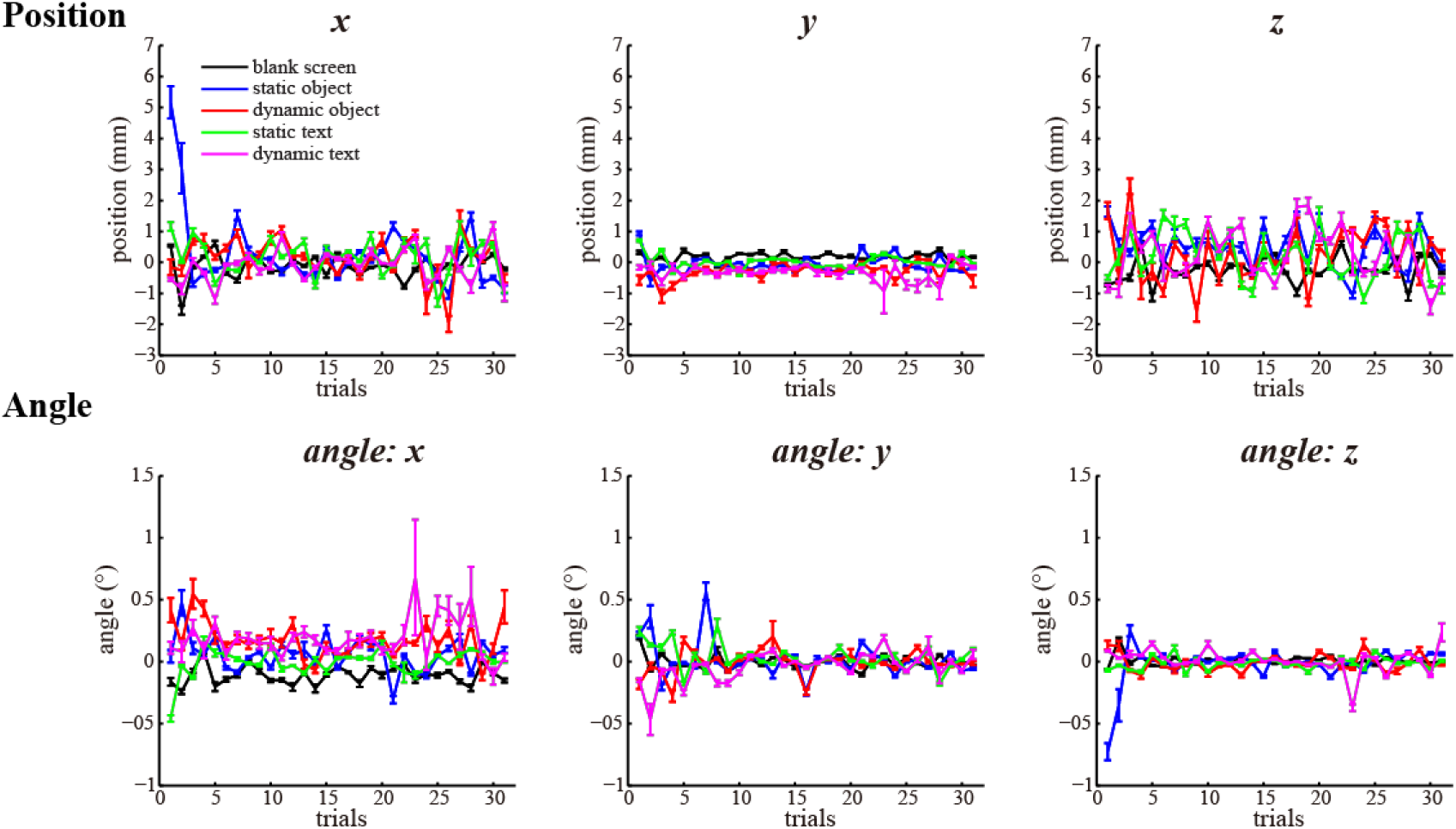
Head movements was plotted as a time of whole experiment - all participants. Head sway traces were shown during the whole experiment including the first practice trial. The mean distance (first row) and angle rotations (second row) were shown for five different visual events. Vertical bars on the columns depict standard error of the mean. Notice that if angle > 0 participant rotates his/her head counterclockwise around axis (from origin to the axis’ positive direction).

We want to quantitatively describe the effect sizes of perturbations. The mean effect was exceptionally small (< 1mm & < 0.5° perturbations, Fig. 7). We analyze the data using one-way ANOVAs with repeated measures for each camera measurement (x, y, z, angle x, angle y, angle z) across the five events (blank screen, static object, dynamic object, static text, dynamic text). A few significant differences emerged in these results. For distance y, there were significant differences among different visual events (F _(4, 105)_ = 3.63, *p* = 0.008). The post hoc tests found participant bend the head more forward in both dynamic object and dynamic text conditions compared to the blank screen condition (p = 0.025 and *p* = 0.021, respectively). For angle x, the differences of visual events were significant (F _(4, 105)_ = 3.54, *p* = 0.009). The post hoc test showed dynamic object induced more head down than that the blank screen (*p* = 0.01). The mean effect, while significant, was very small.

**Fig. 7.**
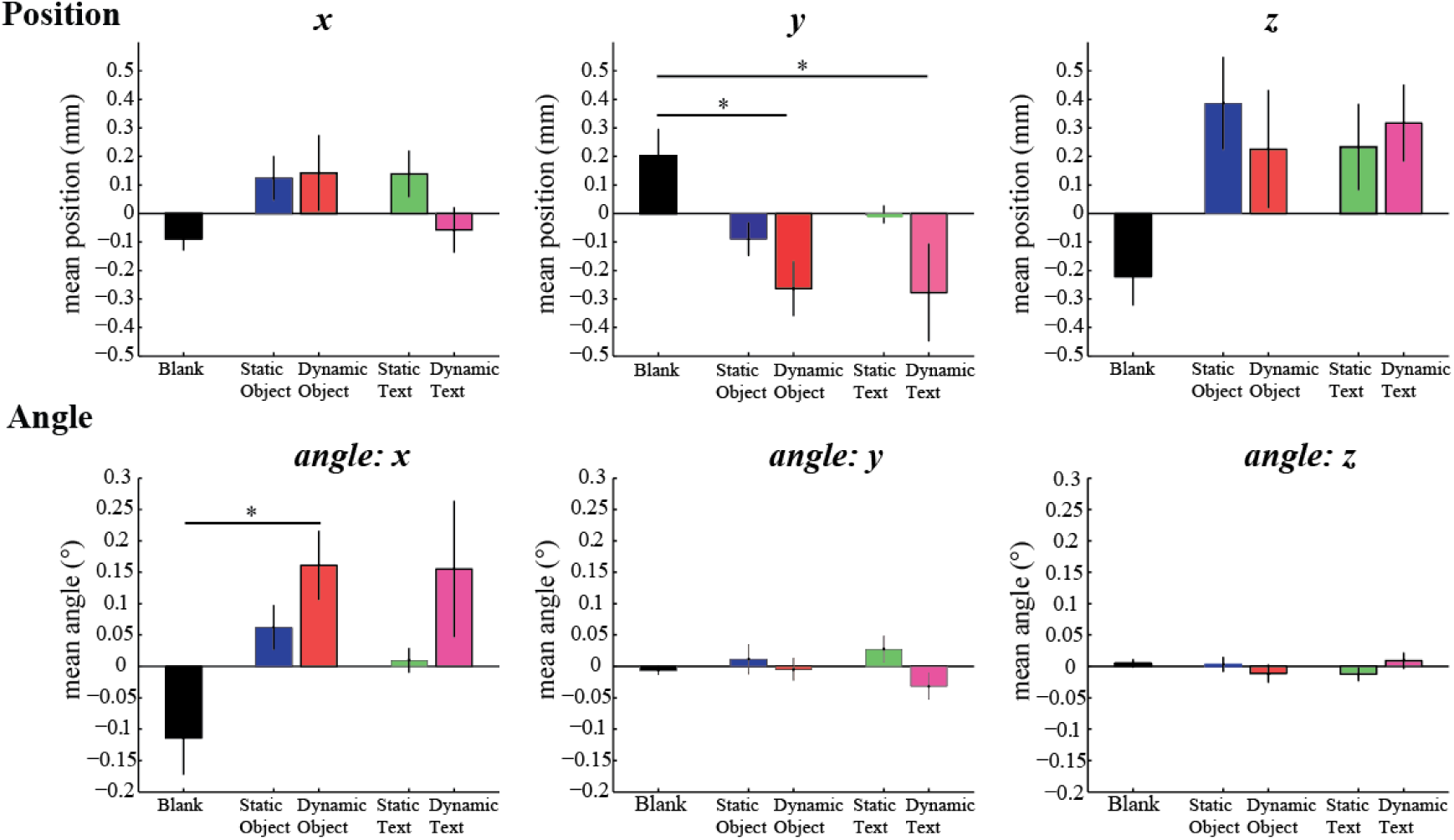
The average distance and angle rotation of head movements– all participants. The distance (first row) and angle rotations (second row) are shown for five different visual events separately. Note that if angle > 0 participant rotates his/her head counterclockwise around axis (from origin to the axis’ positive direction). Vertical bars on the columns depict standard error of the mean. Post hoc differences are marked with * (*p* < 0.05).

The mixed reality environment could affect the variability of head movements within each trial. We calculated the standard deviation (relative to the trial mean) for each trial (Fig. 8). No significant difference was observed by one-way ANOVAs among all comparations. On the whole, the variability of head movement is marginally affected by the stimuli in mixed reality.

**Fig. 8.**
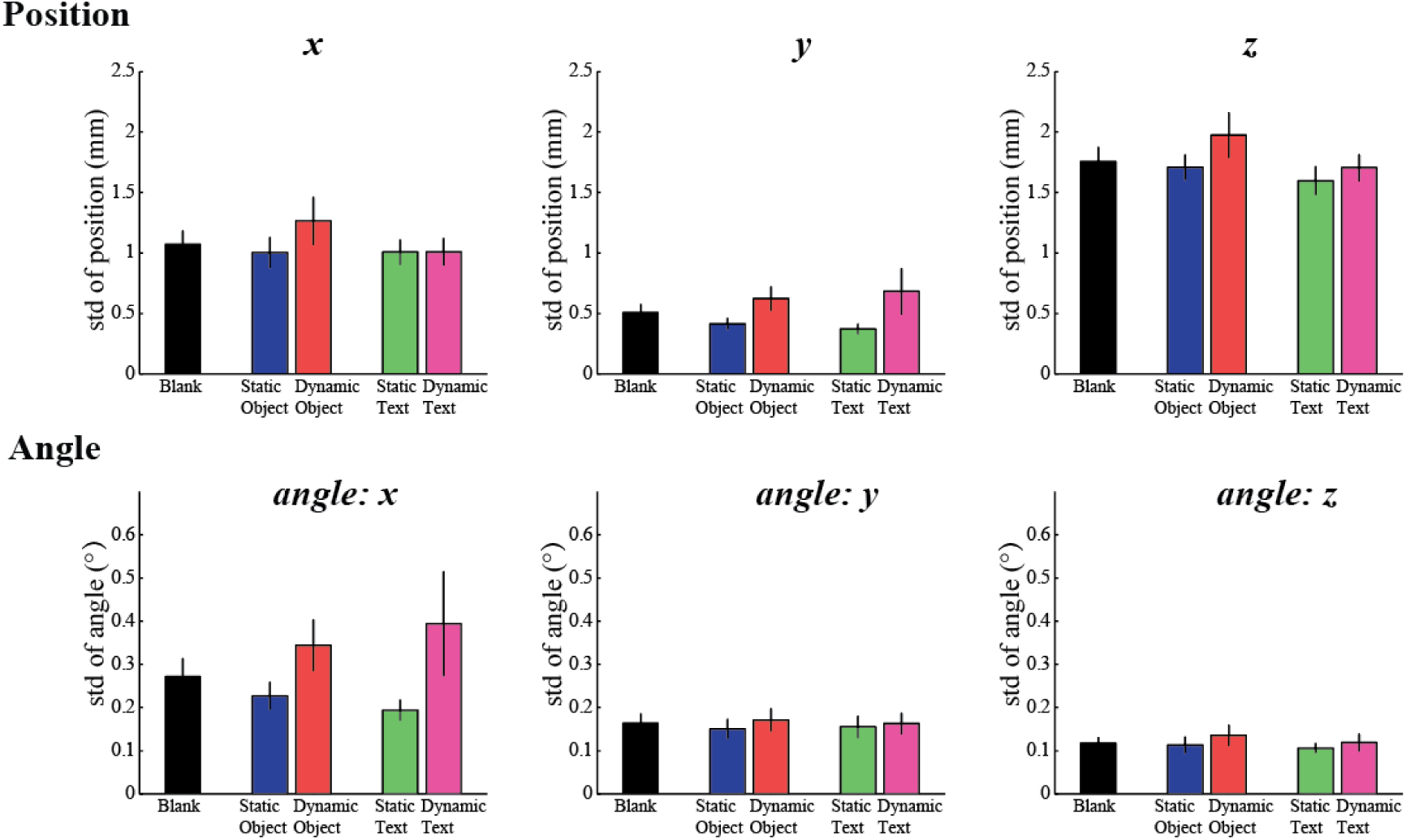
The standard deviation of head movements within each trial – all participants. We calculated the mean head standard deviation for each trial of the same kind visual event. The mean standard deviation of distance (first row) and that of angle rotations (second row) were shown for five different visual events. Vertical bars on the columns depict standard error of the mean of standard deviation.

The mixed reality environment could alternatively affect the variability of head movements across trials. We thus calculated the standard deviation of Root mean square (RMS), across all trials of the same kind visual event (Fig. 9). The standard deviation of RMS values showed that the effect of conditions on head stability was small (< 2.5mm & < 1°). Static conditions tend to have larger variability as compared to dynamic conditions; but this effect is too weak to yield any significant results. The one-way ANOVAs showed no significant differences for the six measurements. Again, the variability of head movement is only marginally affected by visual stimuli presented in the mixed reality.

**Fig. 9.**
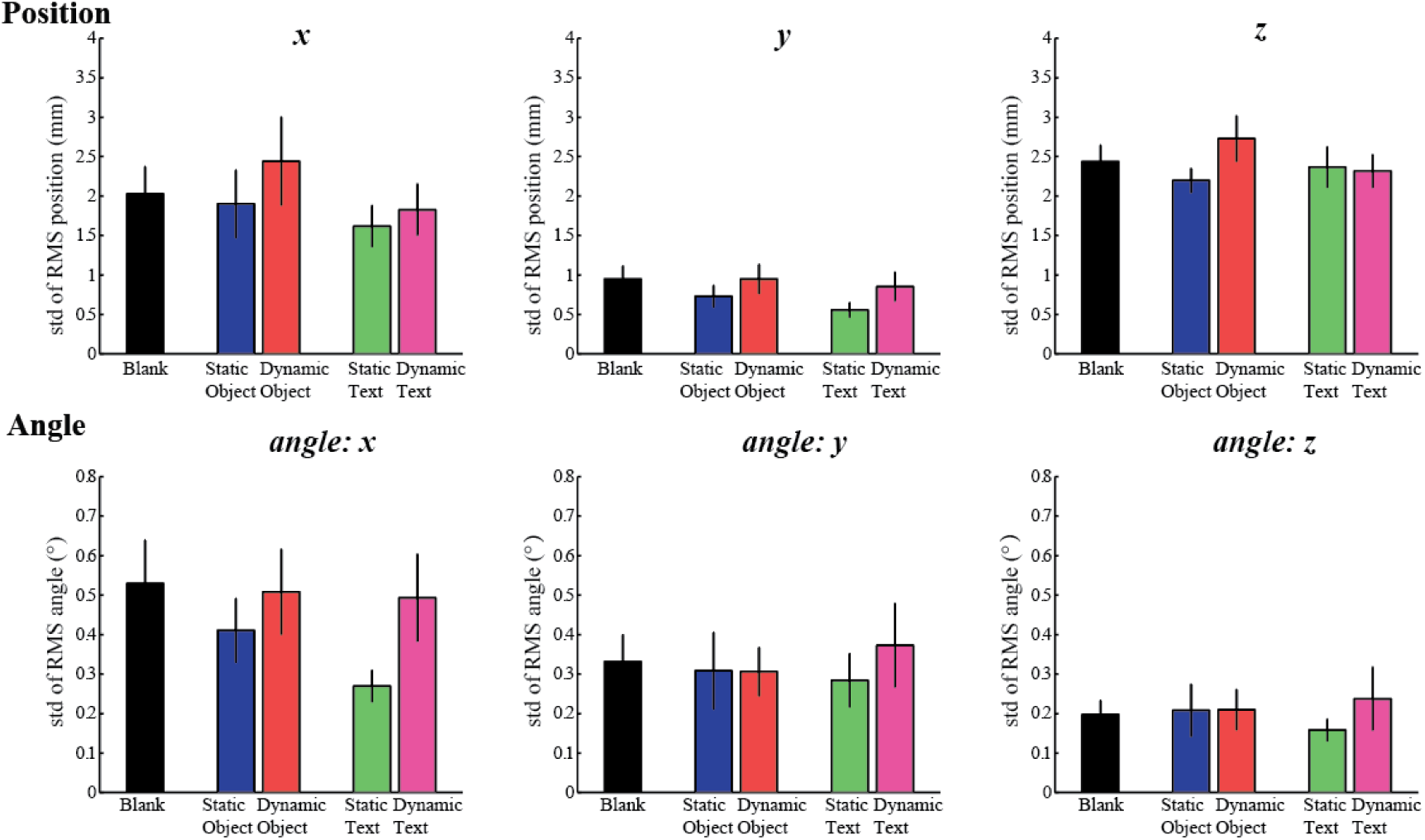
The standard deviation of mean RMS across trials– all participants. The standard deviation of RMS values across trials of distance (first row) and that of angle rotations (second row) are shown for five different visual events. Vertical bars on the columns depict standard error of the mean of standard deviation.

## Discussion

We analyzed the effects of solid object and text presented in Hololens, in both a static and a dynamic setting. The visual stimuli induced some movement, but the effect shown by the mean and variance of head sway, while some significant, was exceptionally small (< 1mm & < 0.5° perturbations). The standard deviation of RMS values also showed that the variability on head sway was small (< 2.5mm & < 1°). The small effects on the presentations become tiny after a few presentations, participants adapted out the effect of mixed reality. Mixed reality does not seem likely to literally knock you off your feet.

We did not investigate a huge range of conditions, just four types of visual events. The duration of the mixed exposure is short in this experiment (< 5 minutes), and each trial lasting 1s is also short, but the duration is enough to show the effect of mixed reality environment. In addition, we did not track the whole body movements, just head movements. One study showed that head movements is stable and reliable as a measure of posture stability[8]. Few studies measured the linear and angular displacements of the head and shoulders simultaneously. However, it hard to move upper body without head movements. We thus think our findings based on head movement can be generalized to other postural stability measures.

The effect of visual events in mixed reality was exceptionally small. One possible reason is that participants still have the peripheral visual inputs which can result in postural compensations [21]. Another possible reason is that mixed reality is different from virtual reality, which completely isolates people from visual reality; this partial connection to reality might help maintain stability. Although we presented dynamic object and text in our experiment, they were simple, repeated visual events; people may adapt to them rapidly across trials. Viewed from this angle, it’s still an open question how much the mixed reality affects people’s quiet stance if we change the visual events to more dramatic ones, for example, making dynamic objects moving faster or appearing from random positions.

We have studied movement behavior while wearing the Hololens. Mixed reality promises to be useful across a broad range of experimental situations. To make it easy for scientists to build on our efforts, we make all our code available online (https://github.com/KordingLab/mixed-reality/tree/master). We hope that this code-base will be useful for movement scientists, psychologists, engineers or practitioners from clinic backgrounds. After all, mixed reality allows presenting visual stimulus to the participant more flexibly than traditional approaches such as using computer monitors or projection screen. This opens the window for a broad range of experimental manipulations and more realistic visual presentations, and it also promises to help scientists investigate more relevant applications to real world questions. We hope our study here serves as an initial endeavor in this direction.

## Acknowledgments

This study was partially supported by National Institutes of Health (NIH 2R01NS063399). The funders had no role in study design, data collection and analysis, decision to publish, or preparation of the manuscript.

## References

1. Horlings CG, Carpenter MG, Küng UM, Honegger F, Wiederhold B, Allum JH. Influence of virtual reality on postural stability during movements of quiet stance. Neuroscience letters. 2009;451(3):227–31.

2. Nishiike S, Okazaki S, Watanabe H, Akizuki H, Imai T, Uno A, et al. The effect of visual-vestibulosomatosensory conflict induced by virtual reality on postural stability in humans. The Journal of Medical Investigation. 2013;60(3.4):236–9.

3. Chiarovano E, de Waele C, MacDougall HG, Rogers SJ, Burgess AM, Curthoys IS. Maintaining balance when looking at a virtual reality three-dimensional display of a field of moving dots or at a virtual reality scene. Frontiers in neurology. 2015;6.

4. Epure P, Gheorghe C, Nissen T, Toader LO, Macovei AN, Nielsen SS, et al. Effect of the Oculus Rift head mounted display on postural stability. International Journal of Child Health and Human Development. 2016;9(3):343.

5. Soffel F, Zank M, Kunz A, editors. Postural stability analysis in virtual reality using the HTC vive. Proceedings of the 22nd ACM Conference on Virtual Reality Software and Technology; 2016: ACM.

6. Robert MT, Ballaz L, Lemay M. The effect of viewing a virtual environment through a head-mounted display on balance. Gait & Posture. 2016;48:261–6.

7. Streepey JW, Kenyon RV, Keshner EA. Field of view and base of support width influence postural responses to visual stimuli during quiet stance. Gait & posture. 2007;25(1):49–55.

8. Kennedy RS, Stanney KM. Postural instability induced by virtual reality exposure: Development of a certification protocol. International Journal of Human-Computer Interaction. 1996;8(1):25–47.

9. Keshner EA, Kenyon RV, Langston J. Postural responses exhibit multisensory dependencies with discordant visual and support surface motion. Journal of Vestibular Research. 2004;14(4):307–19.

10. Horak FB. Postural orientation and equilibrium: what do we need to know about neural control of balance to prevent falls? Age and ageing. 2006;35(suppl 2):ii7–ii11.

11. Peterka R. Sensorimotor integration in human postural control. Journal of neurophysiology. 2002;88(3):1097–118.

12. Oie KS, Kiemel T, Jeka JJ. Multisensory fusion: simultaneous re-weighting of vision and touch for the control of human posture. Cognitive Brain Research. 2002;14(1):164–76.

13. Vuillerme N, Burdet C, Isableu B, Demetz S. The magnitude of the effect of calf muscles fatigue on postural control during bipedal quiet standing with vision depends on the eye–visual target distance. Gait & posture. 2006;24(2):169–72.

14. Duarte M, Zatsiorsky VM. Effects of body lean and visual information on the equilibrium maintenance during stance. Experimental brain research. 2002;146(1):60–9.

15. Vuillerme N, Pinsault N, Vaillant J. Postural control during quiet standing following cervical muscular fatigue: effects of changes in sensory inputs. Neuroscience letters. 2005;378(3):135–9.

16. Wei K, Stevenson IH, Körding KP. The uncertainty associated with visual flow fields and their influence on postural sway: Weber’s law suffices to explain the nonlinearity of vection. Journal of vision. 2010;10(14):4-.

17. Grillon C, Baas JM, Cornwell B, Johnson L. Context conditioning and behavioral avoidance in a virtual reality environment: effect of predictability. Biological psychiatry. 2006;60(7):752–9.

18. Hamm AO, Cuthbert BN, Globisch J, Vaitl D. Fear and the startle reflex: Blink modulation and autonomic response patterns in animal and mutilation fearful subjects. Psychophysiology. 1997;34(1):97–107.

19. Marsh AA, Ambady N, Kleck RE. The effects of fear and anger facial expressions on approach-and avoidance-related behaviors. Emotion. 2005;5(1):119.

20. Smetanin B, Popov K, Kozhina G. Specific and nonspecific visual influences on the stability of the vertical posture in humans. Neurophysiology. 2004;36(1):58–64.

21. Bardy BG, Warren WH, Kay BA. The role of central and peripheral vision in postural control duringwalking. Perception & psychophysics. 1999;61(7):1356–68.

